# Tidal attenuation and poor drainage drive vulnerability of interior microtidal marshes to sea level rise

**DOI:** 10.1101/2023.09.19.558260

**Authors:** Man Qi, Keryn Gedan

## Abstract

Tidal marsh stability under Sea Level Rise (SLR) is driven by feedbacks between marsh macrophyte productivity, sediment accretion, and surface elevation gain. Predictive models and empirical studies often assume a spatially uniform tidal regime across the marsh platform, neglecting hydrological variations caused by tidal attenuation, drainage efficiency and terrestrial groundwater influences. This oversimplification can result in misleading estimates of marsh stability, especially in interior marsh of microtidal systems, which are more vulnerable to SLR-induced deterioration. Here we investigated how spatial variation in hydrology alters marsh vulnerability by comparing groundwater and surface water fluctuations between interior marsh zones and tidal creek edges at two microtidal marshes in the Chesapeake Bay. We assessed the deviation between observed and elevation- and tidally-derived estimates of hydroperiod, daily inundation duration, and soil saturation index (SSI). Using growth-hydroperiod response curves of marsh macrophytes, we assessed how these deviations alter marsh vulnerability predictions. We found that both tidal attenuation and weak drainage play dominant roles in shaping interior marsh hydrology. Due to tidal attenuation, water level fluctuations were muted from a 60 cm range at the marsh edge to a 20 ∼ 30 cm range in marsh interior 500 m away. Therefore, interior marsh exhibits greater hydrological sensitivity to even minor elevation loss, which presents amplified hydrological stress and accelerated vegetation diebacks. Poor drainage in interior marsh result in a persistently higher water level, and slower recession at low tide, leading to prolonged inundation and soil saturation in interior marshes. As a result, assuming a spatially uniform tidal regime underestimated the hydroperiod (by 22-50%) and soil saturation (by 40-50%), and overestimated plant performance (by 10-25%) in interior marsh zones. Our findings highlight the critical need to incorporate tidal attenuation and drainage heterogeneity into empirical studies and numerical simulations to improve marsh resilience prediction and guide adaptive management strategies.

## Introduction

Marsh elevation plays a key role in determining marsh resilience to Sea Level Rise (SLR). Through biomass production, organic matter accumulation and sediment capture, marsh macrophytes regulate surface elevation, forming a dynamic feedback between plant productivity, sediment accretion, and relative sea level [*FitzGerald and Hughes*, 2019; *Gonneea et al.*, 2019; *Kirwan and Mudd*, 2012; *Kirwan et al.*, 2016; *Mudd et al.*, 2010]. Long term marsh stability depends on whether this feedback allows marshes to keep pace with SLR. When marsh elevation exceeds the optimum for primary production, rising water levels stimulate sedimentation and plant growth, facilitating elevation gain. Conversely, if elevation falls below this threshold, productivity declines, sedimentation slows, and the marsh may experience rapid deterioration [*Morris et al.*, 2002].

Because of these elevation-dependent feedbacks, empirical studies and bio-geomorphologic models widely apply the growth response curves of marsh macrophytes to elevation (or water depth at high tide) across entire marsh platforms to estimate marsh stability. Empirical studies extrapolate macrophyte growth responses observed from mesocosm experiments in tidal creeks (i.e. marsh organ experiments) to the entire marsh platforms, assuming hydrological stress is uniform at a given elevation [*Janousek et al.*, 2020; *Janousek et al.*, 2016; *Kirwan and Guntenspergen*, 2012; 2015; *Langley et al.*, 2013; *McLain et al.*, 2020; *Payne et al.*, 2021]. Similarly, bio-geomorphic models predict marsh resilience using a dynamic elevation framework where plant growth is driven by elevation and a spatially uniform tidal range observed in marsh boundary conditions [*D’Alpaos and Marani*, 2016; *Duran et al.*, 2021; *Kirwan and Murray*, 2007; *Kirwan et al.*, 2010; *Marani et al.*, 2013; *Mariotti*, 2016; *Mariotti and Carr*, 2014]. However, these approaches overlook spatial variability in hydrological conditions.

Tidal regimes vary significantly across marsh platforms due to tidal attenuation, drainage conditions, and terrestrial groundwater inputs. As tides propagate inland, energy dissipates and reduces tidal range and water table fluctuations, due to friction, water storage effects, and vegetation resistance [*Lanzoni and Seminara*, 1998; *Möller et al.*, 2014; *Orescanin et al.*, 2019; *Stark et al.*, 2015]. Additionally, poor drainage conditions in interior marsh—regulated by porewater exchange, soil permeability, and tidal creek connectivity—can restrict groundwater recession at low tide, leading to higher baseline water levels and more prolonged inundation (Ursino et al. 2004, Cao et al. 2012a, Xin et al. 2013b, Byers and Chmura 2014, Xin et al. 2022). In some cases, terrestrial groundwater discharges and evapotranspiration play a stronger role than tidal forces in regulating soil moisture in interior marsh [*Guimond et al.*, 2025; *Walker et al.*, 2019; *Xin et al.*, 2013; *Xin et al.*, 2017].

This hydrological heterogeneity challenges the assumption that elevation- and tidally-derived inundation patterns apply uniformly across marsh platforms. Interior marshes often exhibit greater inundation stress and lower resilience than more well-drained marsh edges, contributing to widespread marsh loss in microtidal systems such as the Mississippi Delta, Chesapeake Bay, and Venice Lagoon [*D’Alpaos and Marani*, 2016; *Kearney and Turner*, 2016; *Mitchell et al.*, 2017; *Ortiz et al.*, 2017; *Schepers et al.*, 2017; *Smith et al.*, 2016; *Wang et al.*, 2021]. A recent bio-geomorphic simulation demonstrated that incorporating tidal range dissipation, rather than assuming a uniform tidal range, improves predictions of marsh ponding and degradation [*Zapp and Mariotti*, 2024]. While some studies have acknowledged the limitations of neglecting drainage heterogeneity [*Kirwan and Guntenspergen*, 2015; *McLain et al.*, 2020; *J Morris et al.*, 2013; *Van Putte et al.*, 2022; *Zapp and Mariotti*, 2024], no field-based studies have systematically quantified the different roles that tidal dissipation and drainage condition play in driving interior marsh vulnerability, and how hydrological estimate bias translates into errors in estimating hydrological stress, plant performance, and marsh vulnerability.

To address this gap, we conducted field-based hydrological monitoring at two deteriorating microtidal marsh platforms in the Chesapeake Bay, United States to 1) compare groundwater fluctuations and surface water levels between marsh interior and tidal creeks; 2) quantify the deviation between observed and estimated hydrological metrics, i.e. hydroperiod, inundation duration, and soil saturation; 3) assess the impact of hydrological bias on plant performance and marsh vulnerability using empirical macrophyte growth response curves; and 4) evaluate how muted water table fluctuations in poorly drained interior marsh amplify inundation stress under elevation loss, leading to disproportionate increases in hydroperiod and SSI compared to marsh edges. This study provides empirical evidence that tidal attenuation and drainage constraints drive spatial variability in hydrology. To better predict marsh resilience to SLR, refined monitoring of interior marsh hydrology and improved models that incorporate hydrological heterogeneity would be needed.

### Study area

The Chesapeake Bay is one of the largest microtidal estuaries in the world, and has experienced increasing deterioration in recent decades [*Beckett et al.*, 2016; *Kearney et al.*, 1988]. Within the Bay, we selected two sites, Deal Island (DI) and Farm Creek Marsh (FCM), MD, for study. These two marshes are in the early stages of transition from vegetated marsh to open pond [*Qi et al.*, 2021; *Walker et al.*, 2019]. Areas in various states of deterioration can be found at both sites, but in general, deterioration is more apparent in marsh interiors, where drainage is poor and marsh accretion rates are lower compared with marsh edge [*Qi et al.*, 2021; *Walker et al.*, 2019].

Both sites are brackish marshes, but DI has a higher salinity range (7-15 ppt) than FCM (2-5 ppt) based on measurements in summer of 2019. Emergent vegetation in both marshes is typical of brackish marshes of the region, and includes grasses *Spartina patens*, *Distichlis spicata,* and *Spartina alterniflora*, the sedge *Schoenoplectus americanus*, and the rush *Juncus roemerianus.* The dominant species is *S. alterniflora* in DI, and *S. americanus* in FCM, reflecting differences in salinity between sites and salinity tolerances of these two species. In deteriorating areas of DI, inundation-stressed short-form *S. alterniflora* form a sparse meadow across a spatially homogenous substrate of weak soil strength, whereas in deteriorating areas of FCM, *S. americanus* forms a heterogenous surface of hummocks and hollows.

At the marsh interior of each site, we selected three zones: vegetated, dieback, and pond, to capture the hydrological alteration along the trajectory of marsh deterioration. These marsh zones were within 500 m away from nearest tidal creeks (Fig. 1b and 1c). In DI, we monitored a tidal creek bank located 800 m from interior marsh as a reference point where the water level was likely to resemble an unrestricted tidal fluctuation. We did not instrument such a reference point at FCM because we found that the tidal creek water levels at FCM and DI were highly correlated. Therefore, we took hourly tidal height readings at Bishops Head station, MD from NOAA tide and current website, (Station ID: 8571421, https://tidesandcurrents.noaa.gov/stationhome.html?id=8571421) to represent tidal height at the local scale from FCM to DI. This Bishops Head station is 11 km southeast of FCM, and 12 km Northwest of DI.

**Fig. 1.**
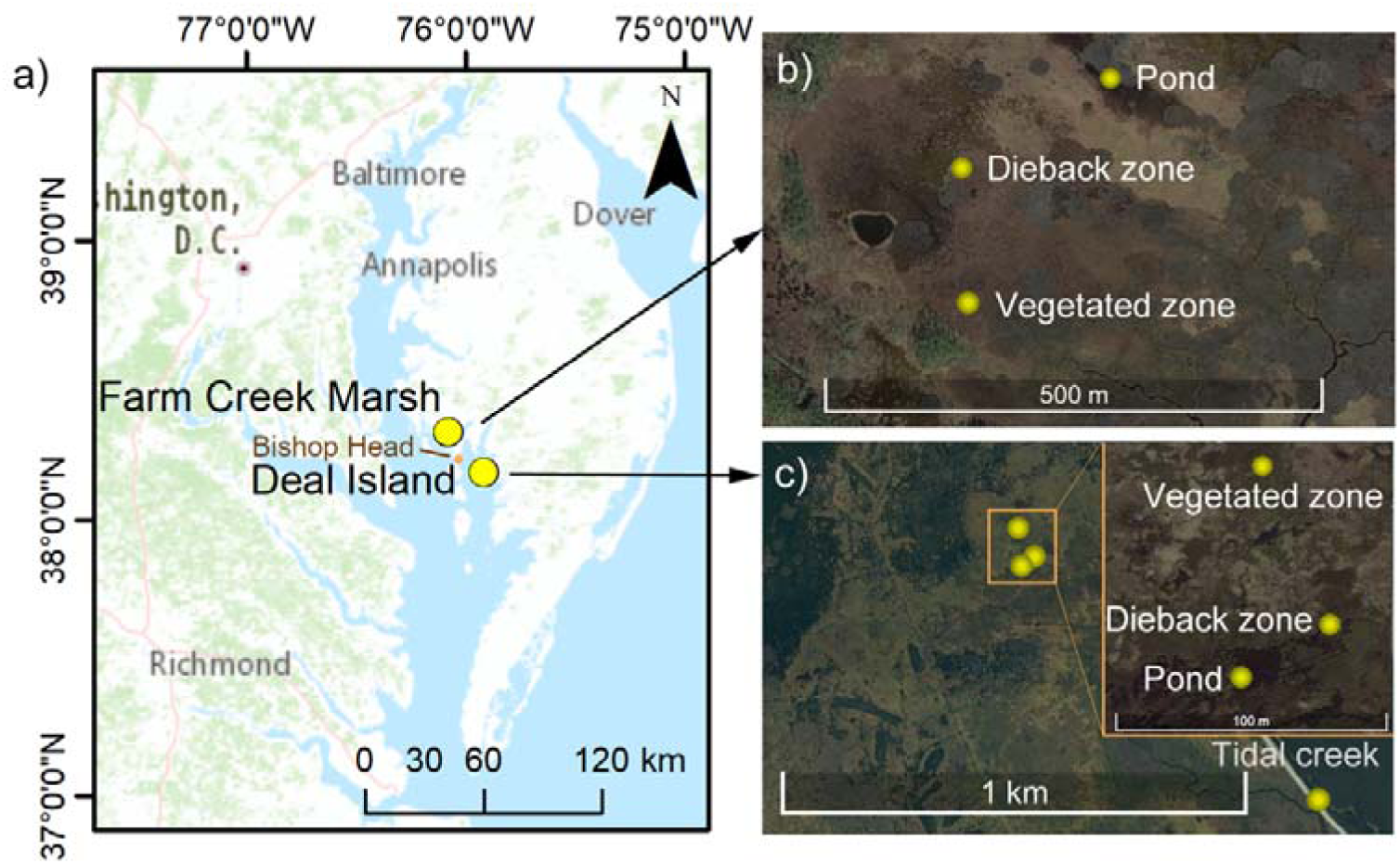
a) Location of the two study sites within the Chesapeake Bay. Shallow wells were located in the vegetated, dieback, and pond zones at each site: b) Farm Creek Marsh (FCM) and c) Deal Island (DI). Tidal height at DI and FCM was acquired from an installed groundwater well within the tidal creek and Bishops Head Tide Gauge, respectively.

## Methodology

### In situ groundwater observation

At each site in April 2019, one groundwater well was installed in each of the vegetated zone, dieback zone, and pond. In DI, another groundwater well was installed in a nearby tidal creek (as shown in Fig. 1) for comparison with the interior part of the marsh. Specifically, at each well site, a one-meter-long well, fully screened other than 10 cm at the top and bottom, was installed to monitor both surface water and groundwater. Hereafter, we use the term water level to refer to water height recorded in the well. The well had a closed, sharp bottom, allowing it to be directly inserted to a depth of 50 cm below the soil surface without the need for packing material. Since installation did not disturb the surrounding soil, flushing was not performed.

Within the wells, water level dataloggers were installed at a depth of 36 ± 6 cm below the soil surface. Water level data was collected at 15 min intervals from May 2019 to March 2020 with HOBO shallow water level loggers (U20L -04). Marsh surface elevation of each zone was measured within a 0.5 m radius of each groundwater well in May 2019 using a Trimble R8 Real Time Kinematic Global Positioning System (RTK-GPS) to reference water table data to the NAVD88 datum and to allow for comparison within and between sites.

### Hydrological metrics

We selected three metrics to represent the hydrological regime of each well location: 1) hydroperiod, 2) daily inundation duration, and 3) soil saturation index (SSI). Hydroperiod was measured as the fraction of time that the marsh was flooded over the observation period (May 2019 to March 2020). Daily inundation duration was calculated as the proportion of time a site remained inundated within a standard 24-hour period (00:00–24:00), based on above continuous water level measurements recorded at 15 min intervals. SSI is the fraction of the tidal period over which the soil remains fully saturated at a given depth, and SSI at depth=0.0 m is equivalent to the hydroperiod [*Xin et al.*, 2010]. We characterized SSI at 0.1 and 0.2 m soil depth.

*In situ* hydrological metrics were calculated from observations of water levels/depth and marsh surface elevation. As a counterpart, estimated hydrological metrics were calculated from tidal height and marsh surface elevation, without employing observational water level data. Comparisons between in situ metrics and estimated metrics were made to estimate the hydrological bias caused by spatially-dependent drainage.

### Influence of hydrological bias on marsh vulnerability

To understand the deviation between *in situ* observations from estimations based solely on elevation and tidal fluctuations (i.e. hydrological bias), and how these affect forecasts of marsh vulnerability to deterioration, we extracted growth response curves of typical macrophytes to predicted and observed hydroperiods. Growth response curves were extracted from the literature, from studies using marsh organ experiments in sites on the Atlantic Coast of the United States. Growth response curves, including the ones used in this study, have been widely used to estimate and forecast marsh vulnerability to sea level rise.

We only adopted growth response curves that used plant biomass (aboveground or belowground) or biomass change relative to inundation duration. Some other response indicators collected in relevant studies included shoot/stem density and stem length, but these were less comparable across studies and could not be related to the hydrological metrics of interest in this study [*Janousek et al.*, 2020; *Kirwan and Guntenspergen*, 2012; *Langley et al.*, 2013].

Based on these criteria, three marsh organ studies contained these variables of interest for four out of five of the typical macrophytes (all except *D. spicata*) at our study sites. Cumulatively, these studies contained 246 measurements of aboveground biomass and 178 measurements of belowground biomass. From these measurements, we normalized aboveground biomass and belowground biomass of a given species from each study separately with the equation of *x_n_*=[*x_i_* - min(*x_i_*)]/[max(*x_i_*)-min(*x_i_*)]. Then we merged the normalized data (with a range of [0∼1]) from different studies to form a unified growth response curve. After a preliminary analysis, we found aboveground and belowground biomass response curves had similar shapes (Appendix S1: Fig. S1), therefore we merged the standardized aboveground biomass and belowground biomass into a uniform biomass response curve to inundation using quadratic regression.

**Table 1.** Data sources of synthesized growth response curves of typical macrophytes to inundation duration.

| Species | Growth response indicators |
| --- | --- |
| *S. americanus* | live aboveground biomass (AGB), live belowground biomass (BGB), root to shoot ratio<sup>a</sup> |
| *S. patens* | live AGB, live BGB, root to shoot ratio<sup>a</sup>; AGB, BGB<sup>b</sup> |
| *S. alterniflora* | AGB , BGB <sup>b</sup>; Total AGB, Live AGB, Total shoot density, End-of-season BGB<sup>c</sup> |
| *J. roemerianus* | total AGB, live AGB, total shoot density, end-of-season BGB<sup>c</sup> |

<sup>a</sup>Kirwan and Guntenspergen [2015]- Blackwater and Transquaking River, Maryland.
<sup>b</sup>Snedden et al. [2015]- Middle Breton Sound and lower Breton Sound, Louisiana.
°Voss *et al.* [2013]- Pine Knoll Shores and Lola, North Carolina.

To estimate the hydrological sensitivity of interior marsh to potential elevation deficiencies as sea level rises, we also analyzed how muted water level fluctuations in interior marsh amplify inundation and saturation compared to tidal creek edges. Taking the marsh platform in DI as an example, we simulated elevation losses of 0 to 0.7 m and compared the resulting hydroperiods and SSIs between interior marsh and tidal creek edges.

## Results

### Marsh elevation and hydrological observations

At both sites, the elevation difference between vegetated zones and ponds was 0.22 m (dashed lines in Fig. 2). However, the spatial distribution of elevation differences varied between the two sites. At DI, elevation differences were gradual, with similar magnitude changes between vegetated, dieback, and ponded zones. In contrast, at FCM, elevation varied minimally between vegetated and dieback zones, but there was a large elevation drop between the dieback zone and ponds.

**Fig. 2.**
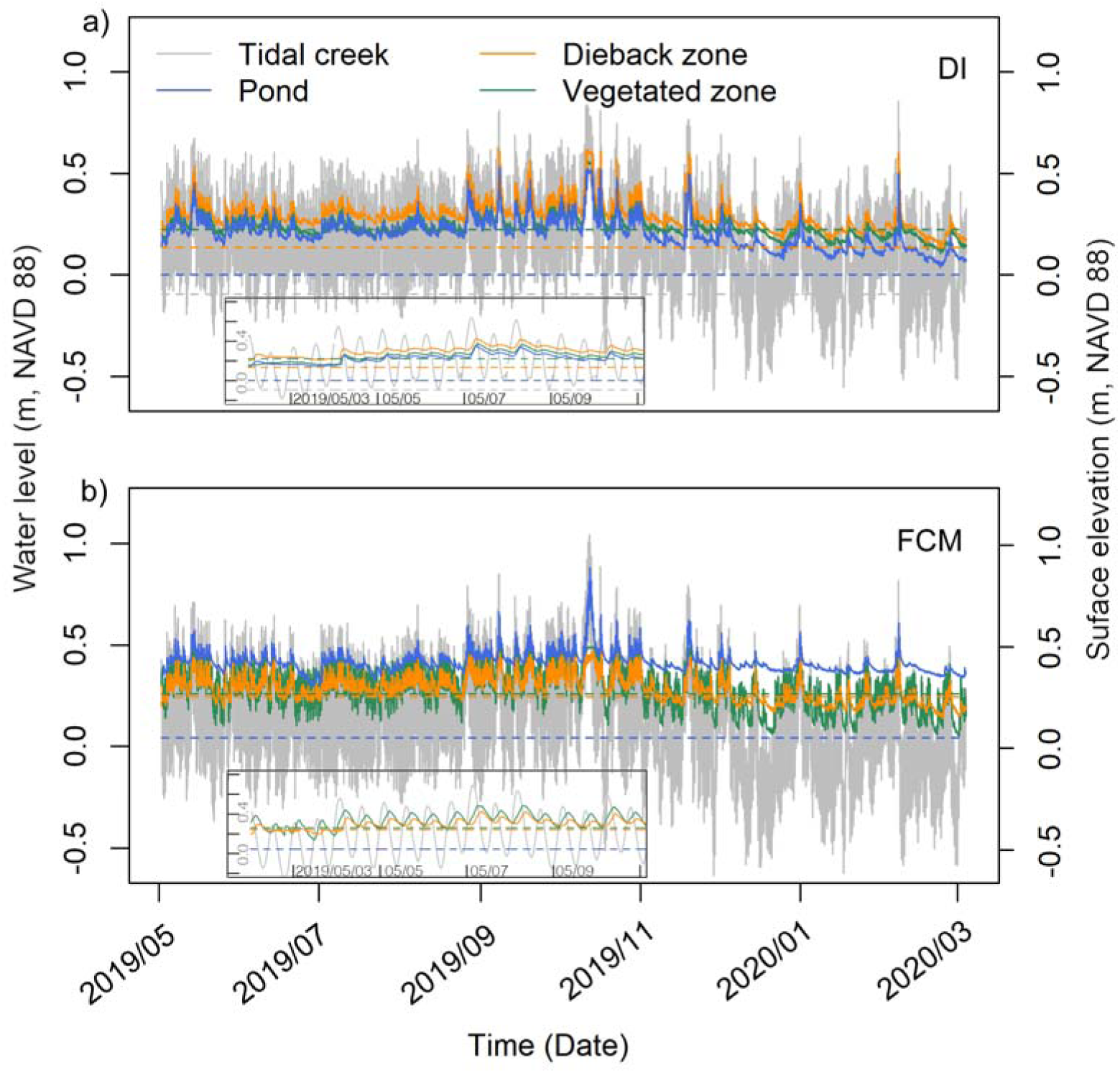
Groundwater table (solid lines) and surface elevation (dashed lines) of different zones (color coded) of deteriorating marshes at DI (a) and FCM (b) (Datum: NAVD88). At DI, ‘tidal creek’ refers to the nearest tidal creek to the interior deteriorating marshes of DI (Fig. 1c), and in FCM, it represents tidal height at the Bishops Head station. Insets in panel a and b shows a zoomed in vision of water level from May 2^nd^ to May 9^th^, 2019.

The water level at DI, monitored at the bank of a tidal creek, exhibited a strong correlation with the tidal level measured at Bishops Head station (Pearson’s correlation coefficient = 0.9605, Fig. 2). The tidal range at both sites was approximately 60 cm (Fig. 3). However, at both sites, water level fluctuations in the interior marsh (vegetated, dieback, and pond zones) were muted to 20 - 30 cm. Additionally, in interior marsh, we observed a consistently higher average water level, 3 - 6 hours lagging in tidal phase, and a slower recession of water level at low tide (insets in Fig. 2).

**Fig. 3.**
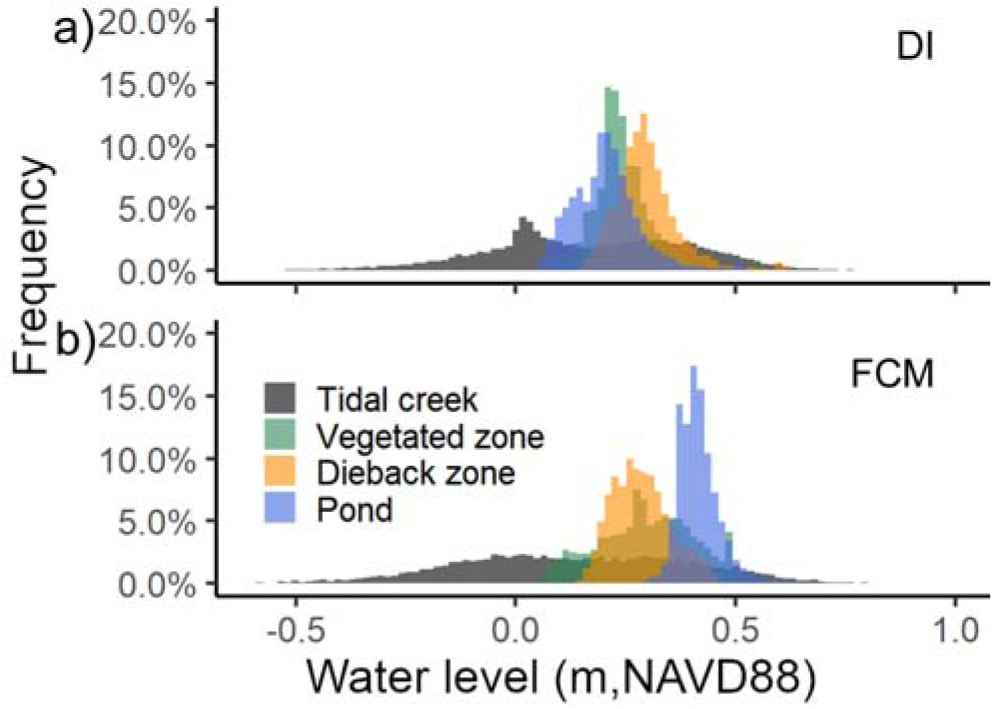
Water level at marsh interiors (vegetated zone, dieback zone, pond) was distributed within a narrow height range compared to tidal creeks at both marsh sites (a-DI; b-FCM). Similar to Fig. 2, ‘tidal creek’ in refers to nearby tidal creek in DI (a), and Bishops Head station in FCM (b).

Differences in water level distributions were also evident across interior marsh zones (Fig. 3). While the range of water table fluctuations was similar at both DI and FCM (20-30cm), the daily variability at DI was lower than at FCM, resulting in a narrower distribution of water level values, particularly for vegetated and dieback zones (Fig. 3).

### Hydrological metrics deviation

The difference between *in situ* observed and estimated hydroperiod (i.e. SSI depth = 0) and SSI at 0.1 and 0.2 m depth was small at the tidal creek but increased within interior marsh (Fig. 4). Estimated hydroperiod and SSI increased linearly with decreasing marsh surface elevation, while observed hydroperiod and SSI increased nonlinearly with increasing elevation and reached a maximum of 100% in most zones at 0.1 m depth and all zones at 0.2 m depth. Soil saturation index at depth was higher than fractional hydroperiod except where both reach a maximum of 100%. The difference between estimated and observed hydroperiod was 22 to 33% within the pond and vegetated zones and 50% in dieback zones. Observed SSI at 0.1 m depth in the dieback zone presented the greatest deviation (50%) from estimated SSI. At 0.2 m depth, the greatest deviation (40%) from estimated occurred in the vegetated zones. This indicates that elevation- tidally- derived estimation consistently underrepresents inundation and soil saturation in poorly drained interior marsh.

**Fig. 4.**
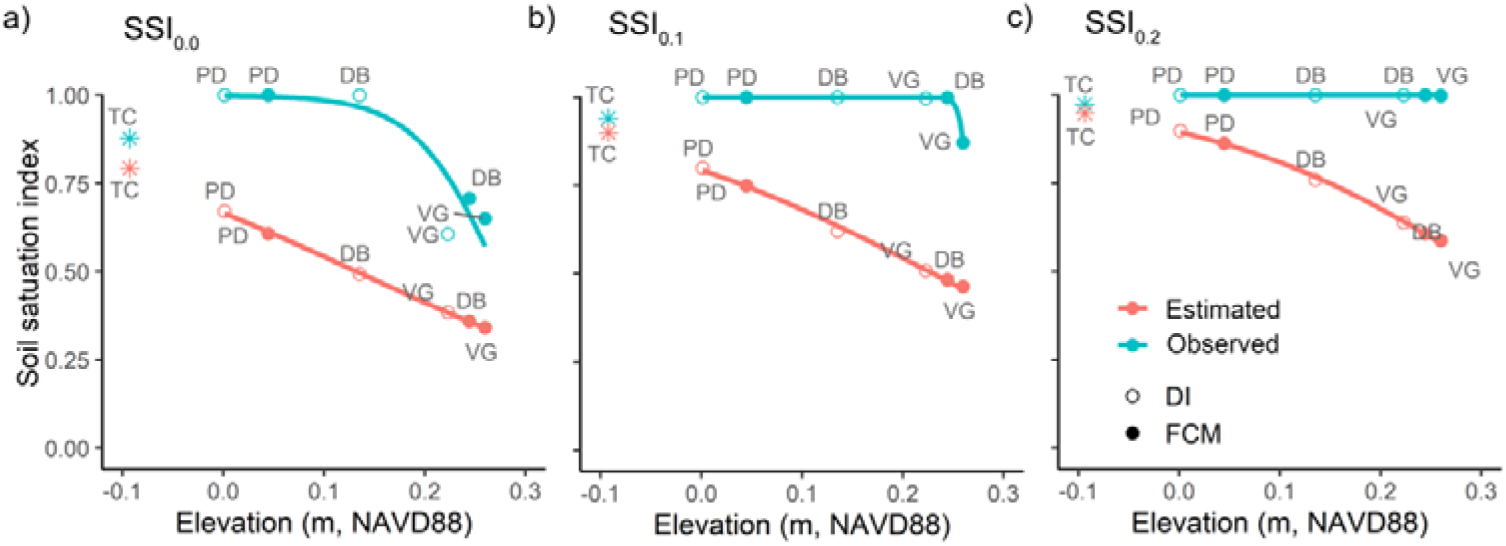
Comparison of soil saturation index (SSI) at soil depth of 0 m (a, SSI_0.0_, or hydroperiod), 0.1 m (b, SSI_0.1_) and 0.2 m (c, SSI_0.2_) between observed (blue) and estimated data (red), derived from elevation and tidal height. TC-tidal creek; PD-pond; DB-dieback zone, VG-vegetated zone. Results of tidal creek in DI was marked separately with *. The response curve of hydroperiod and SSI to elevation in interior marsh was fitted with logistic regression. However, due to the limited range of elevations across the various zones monitored in the field, only a portion of the logistic curve is visible in the plotted data.

The distribution of daily inundation duration differed greatly between in situ observations and estimations. Estimated daily inundation durations averaged between 4 and 8 hrs in all zones for both sites, and increased slightly with the degree of interior marsh deterioration (Fig. 5). However, in observations, vegetated marsh zones were continuously inundated for the entire day on most days (65% of days at DI and 30% of days at FCM). Continuous inundation almost never occurred in any zone in the estimation. For dieback zones and ponds, continuous inundation occurred on 100% of days at DI, and 75% and 100% of days, respectively, at FCM.

**Fig. 5.**
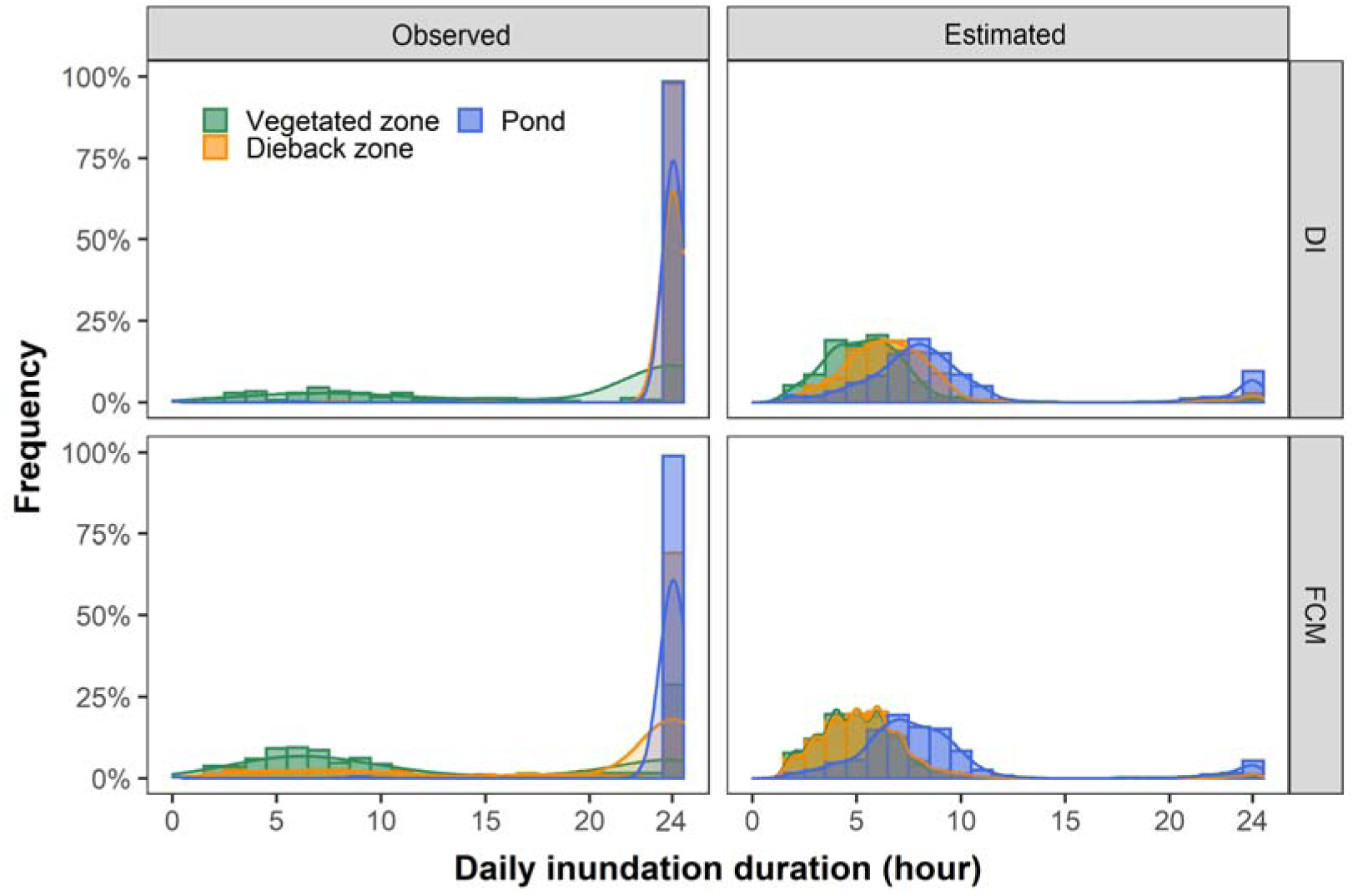
Comparison of the distribution of daily inundation duration between observations and estimations from elevation and tidal height.

### Bias in marsh vulnerability estimation

Among the four macrophytes, only *S. americanus* showed a unimodal growth curve that peaked at intermediate daily durations of inundation; the other three species’ performance gradually decreased with increasing hydroperiod. For vegetated zones of the marsh interior, the estimated hydroperiod (dashed green lines in Fig. 6) is 22%-26% lower than *in situ* observations (solid green lines in Fig. 6). Therefore, the estimation of species performance on the basis of elevation and tidal fluctuations results in an overestimation of plant performance by 10%-25%. This overestimation was most severe for *S. americanus*, the dominant species at FCM. The elevation of the vegetated zone at FCM was located at the optimal level for *S. americanus* performance, according to estimated hydrologic metrics. However, according to observed metrics, the vegetated zone actually experiences suboptimal conditions.

**Fig. 6.**
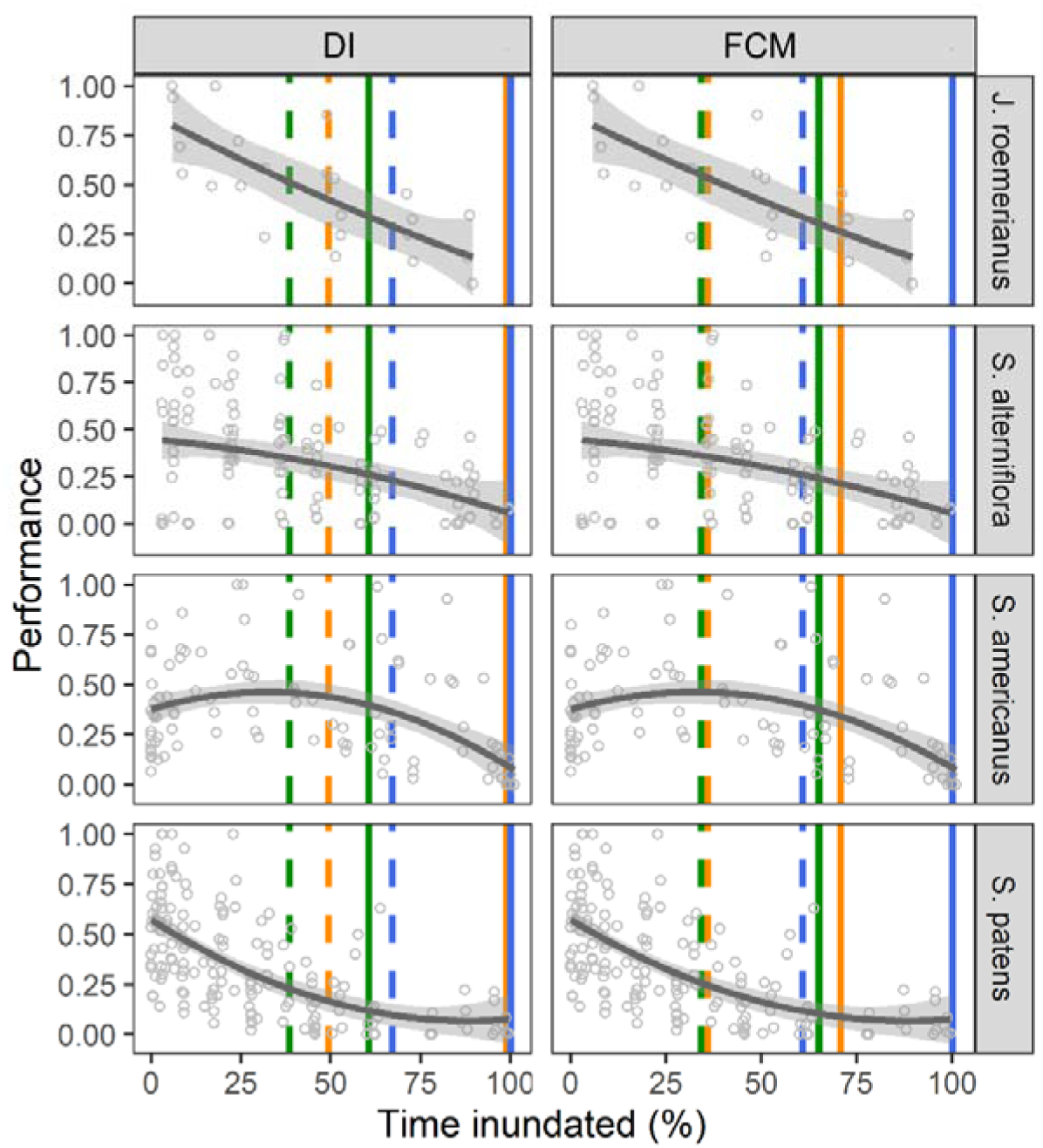
Growth response curves of typical macrophytes to hydroperiod were generated from empirical data [*Kirwan and Guntenspergen*, 2015; *Snedden et al.*, 2015; *Voss et al.*, 2013]. The difference in hydroperiod, estimated using elevation and tidally-based method (dashed lines) and *in situ* observations (solid lines), is displayed for each marsh zone. Colors indicate zone: vegetated (green), dieback (orange), and pond (blue), for Deal Island (DI, left) and Farm Creek Marsh (FCM, right column) sites.

In dieback and pond zones, the estimated hydroperiod indicated a suboptimal but non-lethal status for all species. However, observed hydrologic metrics suggests a hydroperiod at DI that none of the species could survive, which more accurately reflected the vegetation condition. Similar bias also happened in ponds of FCM, but not in dieback zones of FCM, because there was a smaller elevation difference between healthy and dieback areas. Therefore, plant performance and marsh resilience in the vegetated zones of interior marsh was overestimated for all species when elevation and tidally-derived hydrologic regimes were used.

### Difference in sensitivity to elevation deficiency

When elevation drops from 0.36 m to 0.15 m, there is a sharp increase in the hydroperiod in marsh interior (88% increase) relative to the marsh edge (30% increase) (Fig.7). Interestingly, if applying the hydrological sequence in creek bank to interior marsh, the hydroperiod and the SSI in interior marsh can be underestimated at the left side of the intersections of the curves, whereas it is overestimated, albeit to a lesser degree, at the right side of the curve, indicating that at higher elevations of marsh, soils may be more drained and less frequently inundated than expected (Fig. 7). Overall, interior marsh has a narrower elevation range that hydroperiod and soil saturation can quickly increase to be lethal to plants

**Fig. 7.**
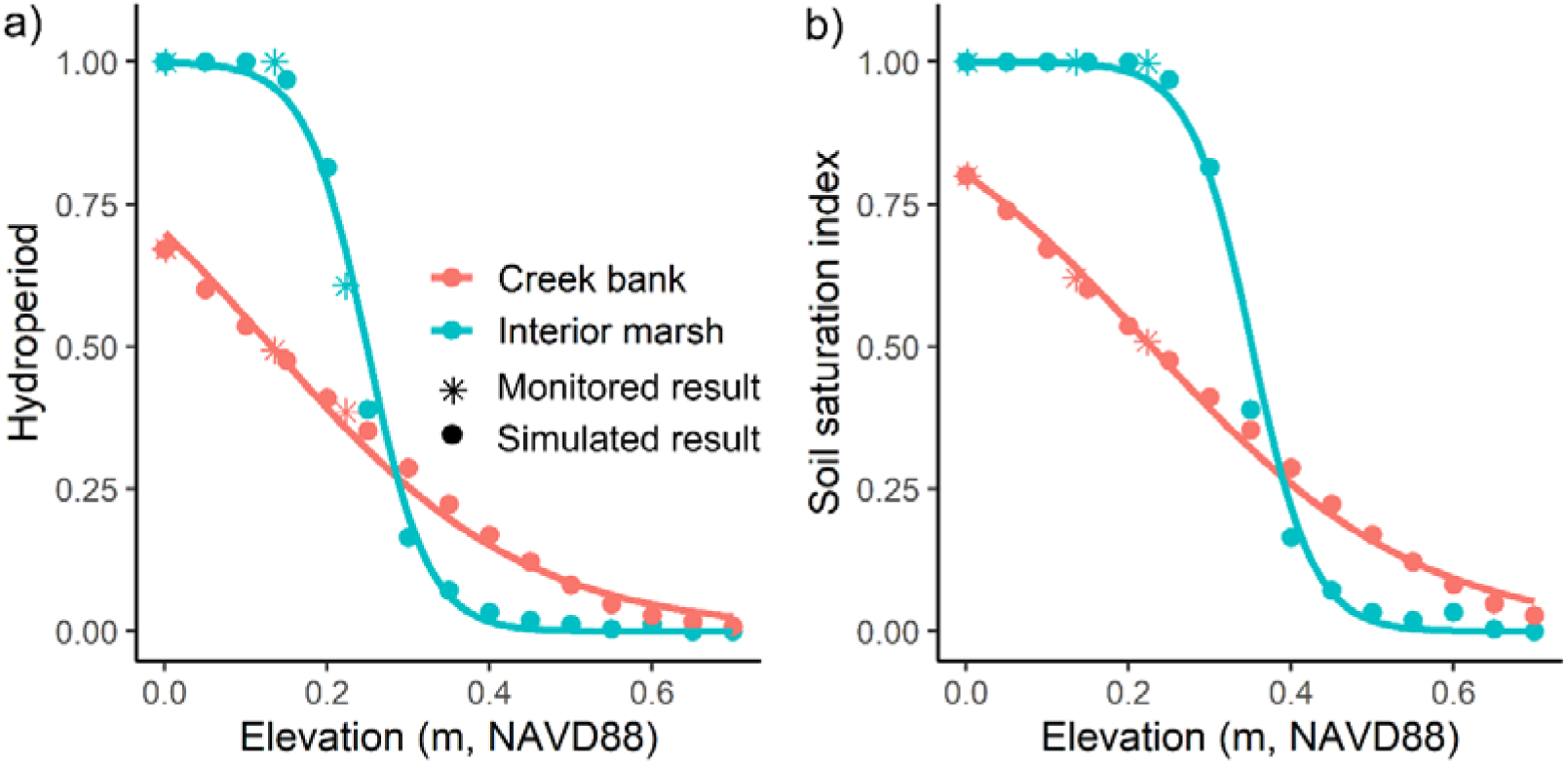
There is a divergent response of hydroperiod (a) and soil saturation index at 0.1 m depth (b) to elevation change in tidal creeks (red) and interior marshes (blue) at Deal Island. Asterisks indicate the observed result in interior marsh, these levels were used to validate the simulated result (filled circles).

## Discussion

Tide station data closely matched tidal creek water levels, confirming strong connectivity between the creek and adjacent marsh edge. In interior marshes, water level fluctuation was muted at both high tide and low tide, indicating strong water attenuation. Additionally, interior marsh maintained a consistently higher average water level, slow recession at low tide and tidal phase lag, suggesting a poor drainage condition in interior marsh. Assuming a uniform tidal regime across the marsh platform lead to an underestimation of hydroperiod by 22-50%, of soil saturation by 40%-50%, and an overestimation of plant performance in interior marsh by 10-25%. Due to the narrow fluctuation in microtidal marshes of the water level, reduced further in interior marsh zones, even minor elevation change can greatly shift the hydroperiod and period of soil saturation. These dramatic deviations in hydrological regimes highlight that spatially heterogenous tidal attenuation and drainage conditions should be accounted for when estimating interior marsh vulnerability to SLR.

### Effect of tidal attenuation and drainage on hydrological regime

We observed four distinct hydrological differences between the marsh edge and interior: 1) tidal fluctuation, 2) tidal phase lag, 3) tidal recession at low tide, and 4) baseline water level. Tidal fluctuation decreased from approximately 60 cm at the marsh edge to 20 ∼ 30 cm at the marsh interior 500 m inland (Fig. 2). This muted fluctuation is caused by energy dissipation as the tide propagates inland, where friction, vegetation resistance, and water storage effects lead to tidal attenuation at both high and low tide peak [*Stark et al.*, 2015; *Xin et al.*, 2022].

Tidal phase lag, slower tidal recession at low tide, and a higher baseline water level in the interior marsh are primarily driven by poor drainage conditions rather than attenuation alone. The interior marsh lags behind the marsh edge tidal dynamics by 3 - 6 hrs (Fig. 2), indicating delayed water exchange between the creek and marsh interior. At low tide, rather than mirroring the rapid water table drop observed at the marsh edge, interior zones retain water for extended periods (Fig. 2). This slow recession of the water table further suggests that drainage inefficiency, rather than tidal attenuation, is the dominant factor controlling interior marsh inundation. In a well-drained system, water would drain efficiently back toward tidal creeks, leading to a more pronounced drop in the eater table at low tide [*Ursino et al.*, 2004]. However, in peatland, the low permeability of peat soils [*Rezanezhad et al.*, 2016] impedes lateral drainage, causing persistent water retention and a higher baseline water table even as tide levels drop (Fig. 2-3). This slower water recession creates prolonged soil saturation and greater hydrological stress on marsh vegetation.

A reduction in water table fluctuations has been observed in other field studies [*Byers and Chmura*, 2014; *M. Cao et al.*, 2012; *Miao Cao et al.*, 2012; *O’Connor and Moffett*, 2015]. However, most observations were collected only over the course of days, and therefore focus on transient asynchronization in groundwater table at different distances from tidal creek. Further, these observed differences have seldom been upscaled to understand or predict differences in vegetation performance and marsh stability between marsh edge and interior. With a year-long record, we found that using estimated values based on elevation and tidal height causes an underestimation of hydroperiod in interior marsh by 25% to 50%, and underestimation of soil saturation by 40% to 50% in the top 20 cm of the soil profile (Fig. 4a).

Through long-term hydrological monitoring, we found that the higher water level in the interior marsh (Fig. 3), caused by poor drainage conditions, contributes to prolonged inundation and soil saturation. Muted water level fluctuations (Fig 2-3), however, increase the interior marsh’s sensitivity to elevation loss (i.e. the threshold response in Fig. 7), making small reductions in elevation lead to disproportionate increases in inundation and soil saturation.

### Spatially heterogeneous marsh vulnerability

In many studies [*D’Alpaos and Marani*, 2016; *Ortiz et al.*, 2017; *Schepers et al.*, 2017], the higher vulnerability of interior marsh is attributed to a lower accretion rate in marsh interiors than at marsh edges [*Kirwan et al.*, 2016]. Our study suggests that the higher inundation at the surface and saturation of the subsurface layer in interior marshes, which is caused by poor drainage, should be a more direct factor. The high frequency of saturation and inundation reduces belowground biomass production, often before aboveground biomass reductions are observed (*Turner et al.* [2004], meaning that these effects can be easily missed in monitoring efforts. This might explain the shallow subsidence rates observed in the Chesapeake Bay, where accretion rates in interior marshes can be much higher than the rate of sea level rise, as high as 9-15 mm/yr, yet marsh surface elevation has decreased 1.8 ± 2.7 mm/yr on average, and this occurred before any evidence of plant dieback was observed [*Beckett et al.*, 2016].

This deficiency of using elevation as the sole indicator of marsh stability, and the importance of drainage in understanding spatial heterogeneity in marsh vulnerability has been reported in prior studies. For example, *Kirwan and Guntenspergen* [2015] reported that *S. americanus* produced 2-4 times greater aboveground biomass in marsh organs than at the equivalent elevations in its natural habitat of interior marsh. Therefore, it is necessary to consider spatial heterogeneity in drainage in future studies of marsh vulnerability and risk.

In addition, inefficient drainage causes overall higher vulnerability of interior marsh than marsh edge, as tidal attenuation increases the sensitivity of hydroperiod to a narrow elevation range in interior marsh. This heightened sensitivity means that deterioration may occur more rapidly and with less warning than at more exterior locations. It may also leave interior marshes less resilient to minor disturbances because a trivial reduction in elevation — approximately 15 cm in this study — can result in a disproportional increase in hydroperiod, dramatically altering growing conditions, triggering plant mortality, or limiting recovery [*Qi et al.*, 2021; *Raposa et al.*, 2017; *Watson et al.*, 2017]. Such elevation deficiency can evolve more quickly in microtidal marshes than meso- and macrotidal marshes. This is because microtidal systems rely more on organic matter production and less on allochthonous sediment inputs [*Ganju et al.*, 2017], therefore minor elevation deficiencies can quickly evolve to be larger through plant-elevation feedbacks.

Deteriorated marsh stages are typically found at low elevation, where peat collapse can follow rapidly after vegetation death [*Day et al.*, 2011]. During the course of marsh deterioration, even minor elevation losses can cause a major change in hydroperiod. Additionally, hydroperiod estimates based solely on elevation and tidal range often fail to capture the true expected performance of plants in these deteriorating microtidal marshes, and may explain the relative lack of anticipation of marsh loss via interior ponding [*Zapp and Mariotti*, 2024]. We conclude that it is essential to account for the heightened baseline water level and reduced water table fluctuations in interior marshes to better predict the elevation threshold and range beyond which runaway marsh collapse is anticipated.

At Deal Island and many other sites, plant displacement in interior marshes has been observed prior to pond formation and expansion [*Qi et al.*, 2021; *Raposa et al.*, 2017; *Watson et al.*, 2017]. Additionally, mortality of flood intolerant species, such as *S. patens* and *D. spicata,* occurred before colonization and replacement by flood tolerant species, such as *S. americanus* and *J. roemerianus* (Fig. 8). The elevation range of *S. americanus* is only a scant 4 to 5 cm lower than *S. patens* [*Holmquist et al.*, 2021], despite a large difference in flood tolerance (Fig. 7, and Qi et al. 2021). It is likely that the steep response curve of hydroperiod and elevation in interior marshes would not provide a sufficient window of opportunity for species displacement. In other words, in the time it takes for die-off of an established plant species, concurrent elevation loss from peat collapse [*Day et al.*, 2011], likely shifts the hydroperiod outside the tolerance range of next most flood-tolerant species, before the latter can establish. Therefore, understanding the elevation building capacity and flood tolerance of different plant assemblages could improve predictions of the elevation trajectory in deteriorating marshes [*Kelleway et al.*, 2017].

**Fig. 8.**
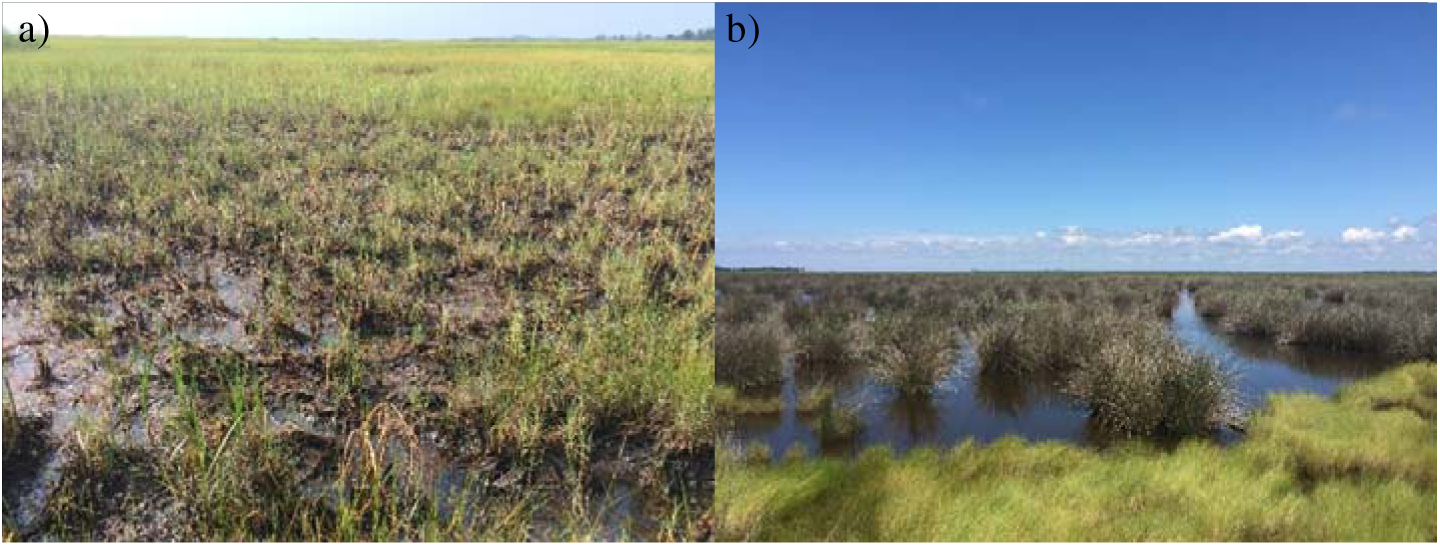
Dieback of *S. patens* and *D. spicata* before being displaced by more flood tolerant species, such as *S. americanus* (a) or *J. roemerianus* (b) in interior marsh of Deal Island.

### Achieving more accurate estimation of interior marsh

The deteriorating marshes selected for this study were located within the range of elevation where elevation and tidally-based estimates severely underestimate hydroperiod and SSI (Fig. 7). This is consistent with observations of rapid vegetation displacement and pond expansion at these same sites over the past decade [*Qi et al.*, 2021; *Taylor et al.*, 2020]. Based on the relationship between observed hydroperiod and elevation, we predict that marsh vulnerability may be overestimated for interior marsh areas at higher elevation or with greater elevation capital. This provides a hypothesis that can be tested in future studies.

Current numeric models that simulate marsh vulnerability to sea level rise assume a spatially uniform hydrological regime (but see *Zapp and Mariotti, 2024*). In general, subsurface flow and its interaction with surface flow are not incorporated in marsh biogeomorphic models [*D’Alpaos and Marani*, 2016; *Duran et al.*, 2021; *Kirwan and Murray*, 2007; *Kirwan et al.*, 2010; *Marani et al.*, 2013; *Mariotti*, 2016; *Mariotti and Carr*, 2014]. Given the dramatic hydrological heterogeneity within marsh platform, we urge future modeling studies to incorporate the impact of tidal attenuation and drainage by applying decreasing tidal range and increasing water level baseline from marsh edge to marsh interior. In this way, the estimates of marsh vulnerability and tools for marsh conservation and restoration can improve.

## Acknowledgement

This study was supported by Washington Biologist’s Field Club (awarded to M Qi). Additional funding was provided by National Natural Science Foundation of China (42306192), Natural Science Foundation award of Hubei Province of China (2023AFB460). We thank Jessica MacGregor, Rose Mohammadi, Richard Peel for helping with the field work. We thank Maryland Department of Natural Resources and Chesapeake Audubon for granting access to field sites.

## References

Beckett, L. H., A. H. Baldwin, and M. S. Kearney (2016), Tidal marshes across a Chesapeake Bay subestuary are not keeping up with Sea-Level Rise, PLOS ONE, 11(7), e0159753.

Byers, S. E., and G. L. Chmura (2014), Observations on Shallow Subsurface Hydrology at Bay of Fundy Macrotidal Salt Marshes, Journal of Coastal Research, 30(5), 1006–1016.

Cao, M., P. Xin, G. Q. Jin, and L. Li (2012), A field study on groundwater dynamics in a salt marsh - Chongming Dongtan wetland, Ecological Engineering, 40, 61–69.

Cao, M., P. Xin, G. Jin, and L. Li (2012), A field study on groundwater dynamics in a salt marsh - Chongming Dongtan wetland, in Ecological Engineering, edited, pp. 61-69.

D’Alpaos, A., and M. Marani (2016), Reading the signatures of biologic–geomorphic feedbacks in salt-marsh landscapes, Advances in Water Resources, 93, 265–275.

Day, J. W., G. P. Kemp, D. J. Reed, D. R. Cahoon, R. M. Boumans, J. M. Suhayda, and R. Gambrell (2011), Vegetation death and rapid loss of surface elevation in two contrasting Mississippi delta salt marshes: The role of sedimentation, autocompaction and sea-level rise, Ecological Engineering, 37(2), 229–240.

Duran, V. O., E. R. Herbert, D. J. Coleman, J. D. Himmelstein, and M. L. Kirwan (2021), Onset of runaway fragmentation of salt marshes, One Earth, 4(4), 506–516.

FitzGerald, D. M., and Z. Hughes (2019), Marsh Processes and Their Response to Climate Change and Sea-Level Rise, Annual Review of Earth and Planetary Sciences, 47(1), 481–517.

Ganju, N. K., Z. Defne, M. L. Kirwan, S. Fagherazzi, A. D’Alpaos, and L. Carniello (2017), Spatially integrative metrics reveal hidden vulnerability of microtidal salt marshes, Nature Communications, 8, 14156.

Gonneea, M. E., C. V. Maio, K. D. Kroeger, A. D. Hawkes, J. Mora, R. Sullivan, S. Madsen, R. M. Buzard, N. Cahill, and J. P. Donnelly (2019), Salt marsh ecosystem restructuring enhances elevation resilience and carbon storage during accelerating relative sea-level rise, Estuarine, Coastal and Shelf Science, 217, 56–68.

Guimond, J. A., et al. (2025), The hidden influence of terrestrial groundwater on salt marsh function and resilience, Nature Water, 3(2), 157–166.

Holmquist, J. R., L. Schile-Beers, K. Buffington, M. Lu, T. J. Mozdzer, J. Riera, D. E. Weller, M. Williams, and J. P. Megonigal (2021), Scalability and performance tradeoffs in quantifying relationships between elevation and tidal wetland plant communities, Marine Ecology Progress Series, 666, 57–72.

Janousek, C. N., B. D. Dugger, B. M. Drucker, and K. M. Thorne (2020), Salinity and inundation effects on productivity of brackish tidal marsh plants in the San Francisco Bay-Delta Estuary, Hydrobiologia, 847(20), 4311–4323.

Janousek, C. N., K. J. Buffington, K. M. Thorne, G. R. Guntenspergen, J. Y. Takekawa, and B. D. Dugger (2016), Potential effects of sea-level rise on plant productivity: species-specific responses in northeast Pacific tidal marshes, Marine Ecology Progress Series, 548, 111–125.

Kearney, M. S., and R. E. Turner (2016), Microtidal Marshes: Can These Widespread and Fragile Marshes Survive Increasing Climate–Sea Level Variability and Human Action?, Journal of Coastal Research, 686-699.

Kearney, M. S., R. E. Grace, and J. C. Stevenson (1988), Marsh Loss in Nanticoke Estuary, Chesapeake Bay, Geographical Review, 78(2), 205–220.

Kelleway, J. J., N. Saintilan, P. I. Macreadie, J. A. Baldock, and P. J. Ralph (2017), Sediment and carbon deposition vary among vegetation assemblages in a coastal salt marsh, Biogeosciences, 14(16), 3763–3779.

Kirwan, M. L., and A. B. Murray (2007), A coupled geomorphic and ecological model of tidal marsh evolution, Proceedings of the National Academy of Sciences, 104(15), 6118–6122.

Kirwan, M. L., and S. M. Mudd (2012), Response of salt-marsh carbon accumulation to climate change, Nature, 489(7417), 550-+.

Kirwan, M. L., and G. R. Guntenspergen (2012), Feedbacks between inundation, root production, and shoot growth in a rapidly submerging brackish marsh, Journal of Ecology, 100(3), 764–770.

Kirwan, M. L., and G. R. Guntenspergen (2015), Response of plant productivity to experimental flooding in a stable and a submerging marsh, Ecosystems, 18(5), 903–913.

Kirwan, M. L., S. Temmerman, E. E. Skeehan, G. R. Guntenspergen, and S. Fagherazzi (2016), Overestimation of marsh vulnerability to sea level rise, Nature Climate Change, 6, 253.

Kirwan, M. L., G. R. Guntenspergen, A. D’Alpaos, J. T. Morris, S. M. Mudd, and S. Temmerman (2010), Limits on the adaptability of coastal marshes to rising sea level, Geophysical Research Letters, 37(23).

Langley, A. J., T. J. Mozdzer, K. A. Shepard, S. B. Hagerty, and J. Patrick Megonigal (2013), Tidal marsh plant responses to elevated CO2, nitrogen fertilization, and sea level rise, Global Change Biology, 19(5), 1495–1503.

Lanzoni, S., and G. Seminara (1998), On tide propagation in convergent estuaries, Journal of Geophysical Research: Oceans, 103(C13), 30793–30812.

Marani, M., C. Da Lio, and A. D’Alpaos (2013), Vegetation engineers marsh morphology through multiple competing stable states, Proceedings of the National Academy of Sciences of the United States of America, 110(9), 3259–3263.

Mariotti, G. (2016), Revisiting salt marsh resilience to sea level rise: Are ponds responsible for permanent land loss?, Journal of Geophysical Research: Earth Surface, 121(7), 1391–1407.

Mariotti, G., and J. Carr (2014), Dual role of salt marsh retreat: Long-term loss and short-term resilience, Water Resources Research, 50(4), 2963–2974.

McLain, N., L. Camargo, C. R. Whitcraft, and J. G. Dillon (2020), Metrics for Evaluating Inundation Impacts on the Decomposer Communities in a Southern California Coastal Salt Marsh, Wetlands, 40(6), 2443–2459.

Mitchell, M., J. Herman, D. M. Bilkovic, and C. Hershner (2017), Marsh persistence under sea-level rise is controlled by multiple, geologically variable stressors, Ecosystem Health and Sustainability, 3(10), 1379888.

Möller, I., et al. (2014), Wave attenuation over coastal salt marshes under storm surge conditions, Nature Geoscience, 7, 727.

Morris, P. V. Sundareshwar, C. T. Nietch, B. Kjerfve, and D. R. Cahoon (2002), Response of coastal wetlands to rising sea level, Ecology, 83(10), 2869–2877.

Morris, J., K. Sundberg, and C. Hopkinson (2013), Salt Marsh Primary Production and Its Responses to Relative Sea Level and Nutrients in Estuaries at Plum Island, Massachusetts, and North Inlet, South Carolina, USA, Oceanography, 26, 78–84.

Mudd, S. M., A. D’Alpaos, and J. T. Morris (2010), How does vegetation affect sedimentation on tidal marshes? Investigating particle capture and hydrodynamic controls on biologically mediated sedimentation, Journal of Geophysical Research: Earth Surface, 115(F3).

O’Connor, M. T., and K. B. Moffett (2015), Groundwater dynamics and surface water-groundwater interactions in a prograding delta island, Louisiana, USA, JOURNAL OF HYDROLOGY, 524, 15–29.

Orescanin, M. M., R. P. Hamilton, and B. Hoffnagle (2019), Tidal choking in an anthropologically modified salt marsh estuary: Improving circulation through constriction removal, Estuarine, Coastal and Shelf Science, 218, 148–162.

Ortiz, A. C., S. Roy, and D. A. Edmonds (2017), Land loss by pond expansion on the Mississippi River Delta Plain, Geophysical Research Letters, 44(8), 3635–3642.

Payne, A. R., D. M. Burdick, G. E. Moore, and C. Wigand (2021), Short-Term Effects of Thin-Layer Sand Placement on Salt Marsh Grasses: A Marsh Organ Field Experiment, Journal of Coastal Research, 37(4), 771–778.

Qi, M., J. MacGregor, and K. Gedan (2021), Biogeomorphic patterns emerge with pond expansion in deteriorating marshes affected by relative sea level rise, Limnology and Oceanography, 66(4), 1036–1049.

Raposa, K. B., R. L. J. Weber, M. C. Ekberg, and W. Ferguson (2017), Vegetation Dynamics in Rhode Island Salt Marshes During a Period of Accelerating Sea Level Rise and Extreme Sea Level Events, Estuaries and Coasts, 40(3), 640–650.

Rezanezhad, F., J. Price, W. Quinton, B. Lennartz, T. Milojevic, and P. Van Cappellen (2016), Structure of peat soils and implications for water storage, flow and solute transport: A review update for geochemists, Chemical Geology, 429.

Schepers, L., M. Kirwan, G. Guntenspergen, and S. Temmerman (2017), Spatio-temporal development of vegetation die-off in a submerging coastal marsh, Limnology and Oceanography, 62(1), 137–150.

Smith, S. M., M. Tyrrell, K. Medeiros, H. Bayley, S. Fox, M. Adams, C. Mejia, A. Dijkstra, S. Janson, and M. Tanis (2016), Hypsometry of Cape Cod Salt Marshes (Massachusetts, U.S.A.) and Predictions of Marsh Vegetation Responses to Sea-Level Rise, Journal of Coastal Research, 33(3), 537–547.

Snedden, G. A., K. Cretini, and B. Patton (2015), Inundation and salinity impacts to above- and belowground productivity in Spartina patens and Spartina alterniflora in the Mississippi River deltaic plain: Implications for using river diversions as restoration tools, Ecological Engineering, 81, 133–139.

Stark, J., T. Oyen, P. Meire, and S. Temmerman (2015), Observations of tidal and storm surge attenuation in a large tidal marsh, Limnology and Oceanography, 60(4), 1371–1381.

Taylor, L., D. Curson, G. M. Verutes, and C. Wilsey (2020), Mapping sea level rise impacts to identify climate change adaptation opportunities in the Chesapeake and Delaware Bays, USA, Wetlands Ecology and Management, 28(3), 527–541.

Turner, R. E., E. M. Swenson, C. S. Milan, J. M. Lee, and T. A. Oswald (2004), Below-ground biomass in healthy and impaired salt marshes, Ecological Research, 19(1), 29–35.

Ursino, N., S. Silvestri, and M. Marani (2004), Subsurface flow and vegetation patterns in tidal environments, Water Resources Research, 40(5).

Van Putte, N., P. Meire, P. Seuntjens, I. Joris, G. Verreydt, L. Hambsch, and S. Temmerman (2022), Solving hindered groundwater dynamics in restored tidal marshes by creek excavation and soil amendments: A model study, Ecological Engineering, 178.

Voss, C. M., R. R. Christian, and J. T. Morris (2013), Marsh macrophyte responses to inundation anticipate impacts of sea-level rise and indicate ongoing drowning of North Carolina marshes, Marine Biology, 160(1), 181–194.

Walker, C. W., T. R. Lester, and C. W. Nealen (2019), Hydrologic study at Farm Creek Marsh, Dorchester County, Maryland, from April 2015 to April 2016 U.S. Geological Survey Scientific Investigations Report, 2019-5032, 12p.

Wang, C., L. Schepers, M. L. Kirwan, E. Belluco, A. D’Alpaos, Q. Wang, S. Yin, and S. Temmerman (2021), Different coastal marsh sites reflect similar topographic conditions under which bare patches and vegetation recovery occur, Earth Surf. Dynam., 9(1), 71–88.

Watson, E. B., K. B. Raposa, J. C. Carey, C. Wigand, and R. S. Warren (2017), Anthropocene Survival of Southern New England’s Salt Marshes, Estuaries and Coasts, 40(3), 617–625.

Xin, P., J. Kong, L. Li, and D. A. Barry (2013), Modelling of groundwater-vegetation interactions in a tidal marsh, Advances in Water Resources, 57, 52–68.

Xin, P., B. Gibbes, L. Li, Z. Song, and D. Lockington (2010), Soil saturation index of salt marshes subjected to spring-neap tides: a new variable for describing marsh soil aeration condition, Hydrological Processes, 24(18), 2564–2577.

Xin, P., T. Z. Zhou, C. H. Lu, C. J. Shen, C. M. Zhang, A. D’Alpaos, and L. Li (2017), Combined effects of tides, evaporation and rainfall on the soil conditions in an intertidal creek-marsh system, Advances in Water Resources, 103, 1–15.

Xin, P., et al. (2022), Surface Water and Groundwater Interactions in Salt Marshes and Their Impact on Plant Ecology and Coastal Biogeochemistry, REVIEWS OF GEOPHYSICS, 60(1).

Zapp, S. M., and G. Mariotti (2024), Tidal dissipation morphodynamic feedback triggers loss of microtidal marshes, Geology, 52(5), 326–330.

