## Appendix S1 for "Tidal attenuation and poor drainage drive vulnerability of interior microtidal marshes to sea level rise"

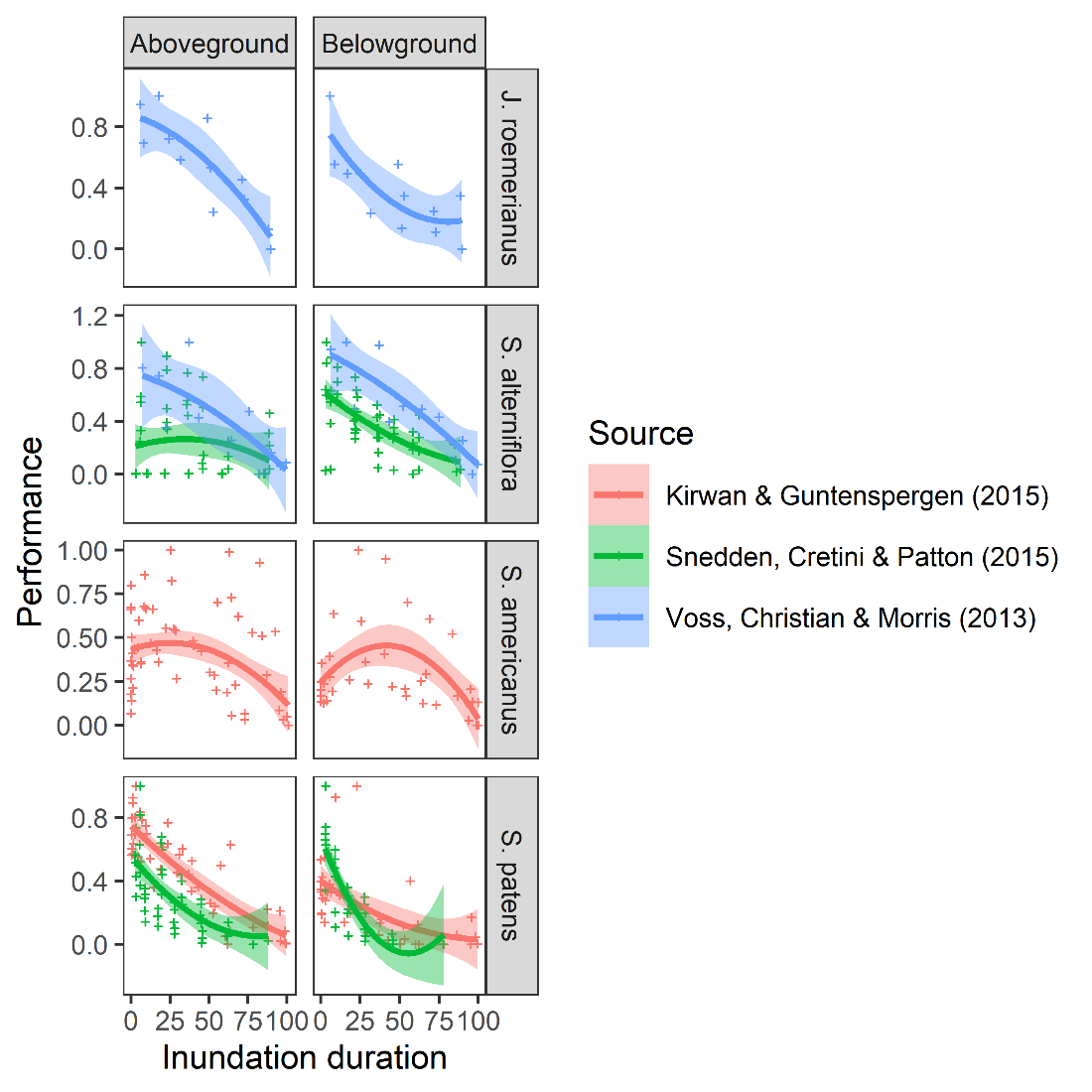

Fig. S1 Respond of aboveground and belowground biomass to increasing inundation duration.
